# Continuing decline of the eastern quoll in Tasmania

**DOI:** 10.1101/2022.03.10.483855

**Authors:** Calum X Cunningham, Zach Aandahl, Menna E Jones, Rowena Hamer, Christopher N Johnson

## Abstract

Like many other Australian mammals, the eastern quoll (*Dasyurus viverrinus)* was widespread on the Australian mainland but went extinct there during the 20th century. The species remained abundant in Tasmania until a rapid decline occurred from 2001 to 2003, coinciding with a period of unsuitable weather. We provide an updated analysis of eastern quoll population trends in Tasmania by analysing a Tasmania-wide time series of annual spotlight counts (1985-2019). Eastern quolls were widespread and abundant in Tasmania until the early 2000s. A distinct change occurred in the early 2000s in the east and northeast, which led to severe population reductions. However, we present new evidence of an earlier decline in the north (mid-1990s) and a more recent decline around 2009 in the south. Range-wide declines have continued unabated during the last decade, resulting in a ∼67% decline (since the late 1990s) in the area with high quoll abundance. Although the timing of the major decline in the early 2000s coincided with unfavourable weather, the continuing decline and more recent change points suggest other causes are also involved. We can no longer assume that the existence of eastern quolls in Tasmania ensures the species’ long-term survival, highlighting the urgent need to increase efforts to conserve the remaining populations in Tasmania.

## Introduction

Like many small and medium-sized Australian mammal species (Woinarski *et al*. 2015), the eastern quoll (*Dasyurus viverrinus*) was once widespread across the south-eastern Australian mainland (Peacock & Abbott 2014) but has declined severely since European colonisation. Eastern quolls went extinct on the Australian mainland in the 20^th^ century, with the last confirmed sighting in 1963, although the species possibly survived at very low abundance for some time after (Frankham *et al*. 2017). Recently, a population was reintroduced to Booderee National Park, NSW, as a first step in re-establishing wild eastern quoll populations on the Australian mainland (Robinson *et al*. 2020).

In the absence of red foxes (*Vulpes vulpes*), eastern quolls remained abundant and widespread in Tasmania until the early 2000s when densities declined sharply (Fancourt *et al*. 2013; Fancourt *et al*. 2015a). While these population declines were initially attributed to the mesopredator release of feral cats, stemming from the disease-induced decline of the Tasmanian devil (Hollings *et al*. 2014), later research suggested they were associated with a period of unsuitable weather over much of the species’ distribution from 2001-2003 (Fancourt *et al*. 2015a). Following a return to suitable weather, the population did not recover, suggesting other factors might be preventing recovery, such as a “predator pit” caused by feral cats (Fancourt *et al*. 2015a). Based on a 52% decline of spotlighting sightings in the decade to 2009, eastern quolls were listed as Endangered under the IUCN Red List (Burbidge & Woinarski 2016). The species is listed as Endangered under the federal *Environment Protection and Biodiversity Conservation Act 1999* but is not listed under the Tasmanian *Threatened Species Protection Act 1995*.

The most recent published assessment of eastern quoll population trends used survey data until 2009 (Fancourt *et al*. 2013; Fancourt *et al*. 2015a). The aim of this paper is to provide an updated analysis of eastern quoll population trends in Tasmania, review possible causes of decline, and suggest actions for recovery. We use a 35-year (1985-2019) standardised spotlighting dataset from across Tasmania to characterise changes in the distribution and relative abundance of eastern quolls. We apply recent advances in species distribution modelling to fit spatiotemporal models of relative abundance through time. We also use change-point models to identify the timing of change in regional population trajectories and evaluate whether the onset of decline is consistent with the weather-induced decline hypothesis.

## Materials and Methods

### i) Long-term spotlighting data

The Tasmanian State Government has conducted standardised annual spotlighting surveys across Tasmania from 1985-2019, totalling 5,761 transect counts (Fig 1). The spotlight surveys were initially established to monitor populations of herbivores subject to harvest, while recording all sightings of non-domestic mammals, including eastern quolls (Hocking & Driessen 1992). The same transects are surveyed each year, however the number of transects increased from approximately 130 in the 1980s to approximately 170 in the 1990s (Supplementary Table S1). We visually evaluated the effect of survey additions in the 1990s, which generally did not appear to contribute to the observed trends (Fig S1). The most notable difference occurred in the ‘South’ region, where the number of transects increased from 9 in the 1980s to 28 in 1992 (Table S1). In this region, analyses of the full dataset resulted in more uniform average annual detections and weaker inter-annual trends. We therefore opted to analyse the full dataset as the more conservative option and because it comprised a more comprehensive sample of the region.

**Figure 1:**
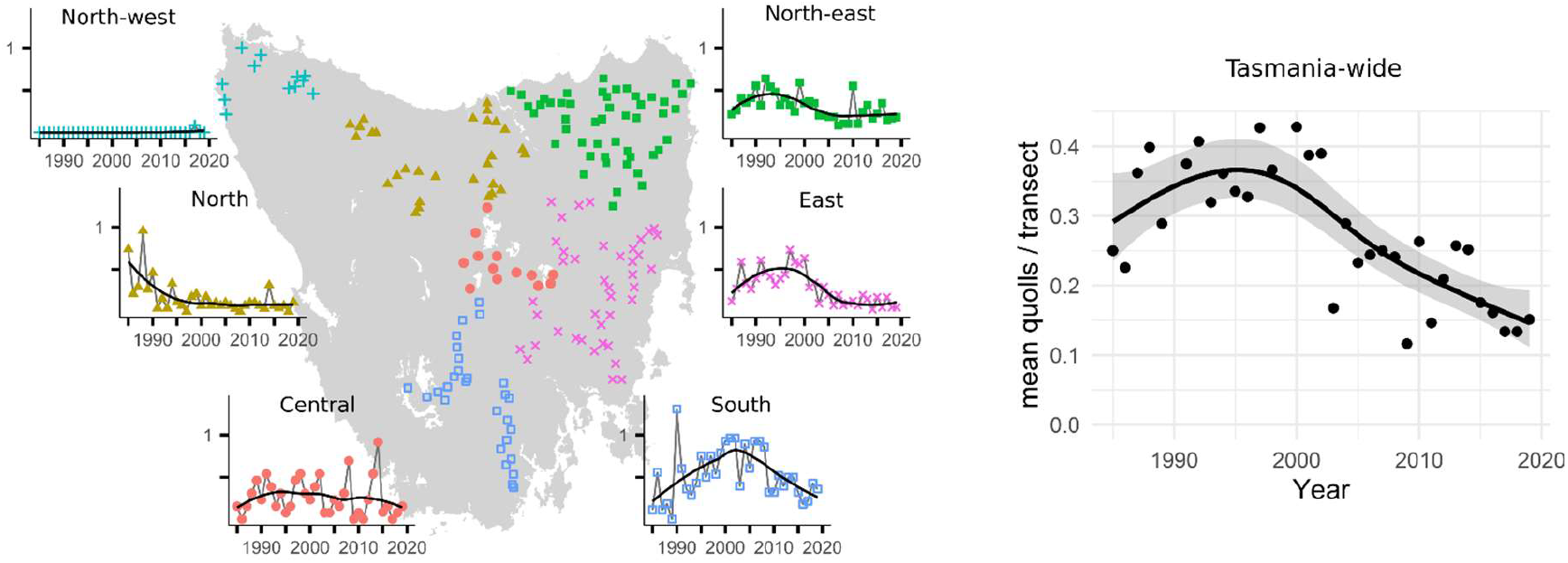
Trends of eastern quoll detections in the long-term spotlighting data. On the left, transects are grouped into IBRA bioregions (IBRA DSEWPC 2013). Data points in the graphs show the mean number of eastern quoll detections on all transects in a bioregion. The black lines are a simple loess smoother. The right graph shows the mean number of quolls sighted per transect, across all of Tasmania.

Each transect follows a 10-km strip of hardened or gravel road. Transects are driven once per year at a speed of 20 km/hr, with one person using a handheld spotlight to observe animals on both sides of the road while a second person records animal sightings (for details, see Hocking & Driessen 1992; Hollings *et al*. 2014). Transects are surveyed once each year during the summer months to ensure comparability across years. However, because transects are only surveyed once per year, this prevents the use of statistical techniques that separate the state and detection processes. While we consider the count of eastern quolls per transect as an index of population density (see Fig 1 for trends), we acknowledge that the spotlight counts are a combined measure of the detection and abundance processes (MacKenzie & Kendall 2002). Overall, this long-term dataset is of rare spatial and temporal scope, and has been useful to monitor population trends and infer the causes of population dynamics in a wide range of species (Hollings *et al*. 2014; Lazenby *et al*. 2018; Carver *et al*. 2021; Cunningham *et al*. 2021; Cunningham *et al*. 2022).

We grouped transects into six regions (Fig 1) based on proximity. We attempted to align our regions with the IBRA bioregions, with some aggregation necessary in areas where few transects fell in an IBRA bioregion. Our regions align with IBRA bioregions as follows: ‘North-west’ fits within the King bioregion; ‘North’ comprises the Northern Slopes bioregion, with three south-western transects falling just over the boundary of the Central Highlands bioregion; ‘Central’ falls within the eastern half of the Central Highlands bioregion; ‘Northeast’ comprises both the Ben Lomond and Flinders bioregions; ‘East’ comprises the Northern Midlands and South East bioregions; ‘South’ fits entirely within the Southern Ranges bioregion (IBRA DSEWPC 2013).

### ii) Statistical analysis

#### Spatial changes in the distribution of eastern quolls

We modelled changes in the distribution of eastern quolls using a spatiotemporal autoregressive model. This approach aims to characterise the realised distribution of eastern quolls through time. We fitted the models using integrated nested Laplace approximation (INLA), an accurate and fast option for Bayesian inference from spatial data. To fit the models, we used the ‘inlabru’ R package (Bachl *et al*. 2019; R Core Team 2019), which is an extension of the R-INLA package (Rue *et al*. 2009; Bakka *et al*. 2018).

INLA is useful for modelling spatial data because spatial dependence between observations can be modelled using a continuous correlation process known as a Gaussian random field. A Gaussian random field is a spatially continuous process where random variables at any point in space are normally distributed and are spatially correlated with other random points via a continuous correlation process (Bachl et al. 2019). R-INLA approximates the continuous correlation process using a stochastic partial differential equation (SPDE). The SPDE requires discretising the study domain into a series of abutting triangles, known as a mesh. We discretised Tasmania into a mesh with internal edge lengths of 10 km, corresponding to the scale of the spotlight transects (inla.mesh.2d function of the R-INLA package).

Using the mesh, a Gaussian random field is then approximated by an SPDE. We constructed a spatiotemporal SPDE to allow the spatial correlation between observations to change through time. To model the temporal dependence in eastern quoll observations between consecutive years, we began by fitting first- and second-order autoregressive models (Gómez-Rubio 2020). These models, however, had convergence problems, probably because of relatively sparse sightings and moderate year-to-year variability. To smooth some of the year-to-year variability, we grouped surveys into five-year periods. We fitted the spatio-temporal random field using a first-order autoregressive model (Gómez-Rubio 2020), which in effect correlated observations in consecutive five-year periods. We used a Matérn correlation structure for the SPDE, and specified a prior probability of 0.1 that the range was less than 10 km and a 0.5 probability that the standard deviation was greater than 1 km. For further details about SPDEs, see Lindgren et al. (2011).

We used the count of eastern quolls per transect as the response variable and fitted the model using the negative binomial distribution to account for overdispersion (Ver Hoef & Boveng 2007). We modelled counts in response to the spatio-temporal Gaussian random field described above. From this model, we produced maps of the relative abundance of eastern quolls in five-year periods from 1985 to 2019. We visualised areas with high relative abundance using a contour around the areas in which the model-estimated relative abundance was at least one quoll per transect.

#### Change-point modelling

We use change point models to infer if, and when, changes in transect counts of eastern quolls occurred in the different regions of Tasmania. We modelled change points by performing segmented regression with the R package ‘mcp’ (Lindeløv 2020). Because the literature contains strong evidence for a step-change in quoll abundance (Fancourt *et al*. 2013; Fancourt *et al*. 2015a), we considered three-simple models: 1) a one-segment null model with no change point, 2) a two-segment model that included a single disjoined change point (a step change), and 3) a two-segment model with a disjoined change point with differing variance between the segments. We fitted these models for each region and selected the best-performing model using the Estimated Log Predictive Density (ELPD) from approximate leave-one-out cross-validation, in which larger values represent better predictive accuracy.

## Results

Spatio-temporal distribution models show that eastern quolls occupied much of eastern Tasmania at high densities until the early 2000s (Fig 2), when the model-estimated area with at least one quoll per spotlight transect peaked at 11,500 km^2^ (Fig 3). The population then went into a decline that has continued for almost two decades (Fig 3). Eastern quolls are now widespread and abundant only in southern Tasmania, along with small and isolated pockets in other areas (Fig 2). Since the population peak in the late 1990s, the Tasmania-wide index of abundance has declined by 60% (Fig 1) and the model-estimated area with at least one quoll per transect has declined by 67%, now at ∼3,750 km^2^(Fig 3). This declining trend has continued in the most recent decade (Fig 1 & Fig 3). Note that these estimates of percentage change are relative to the population peaks in the late 1990s rather than commencement of monitoring in 1985, as there is suggestion that carnivore populations in the 1980s were still recovering from widespread use of strychnine poisoning (Hawkins *et al*. 2006; McCallum & Jones 2006).

**Figure 2:**
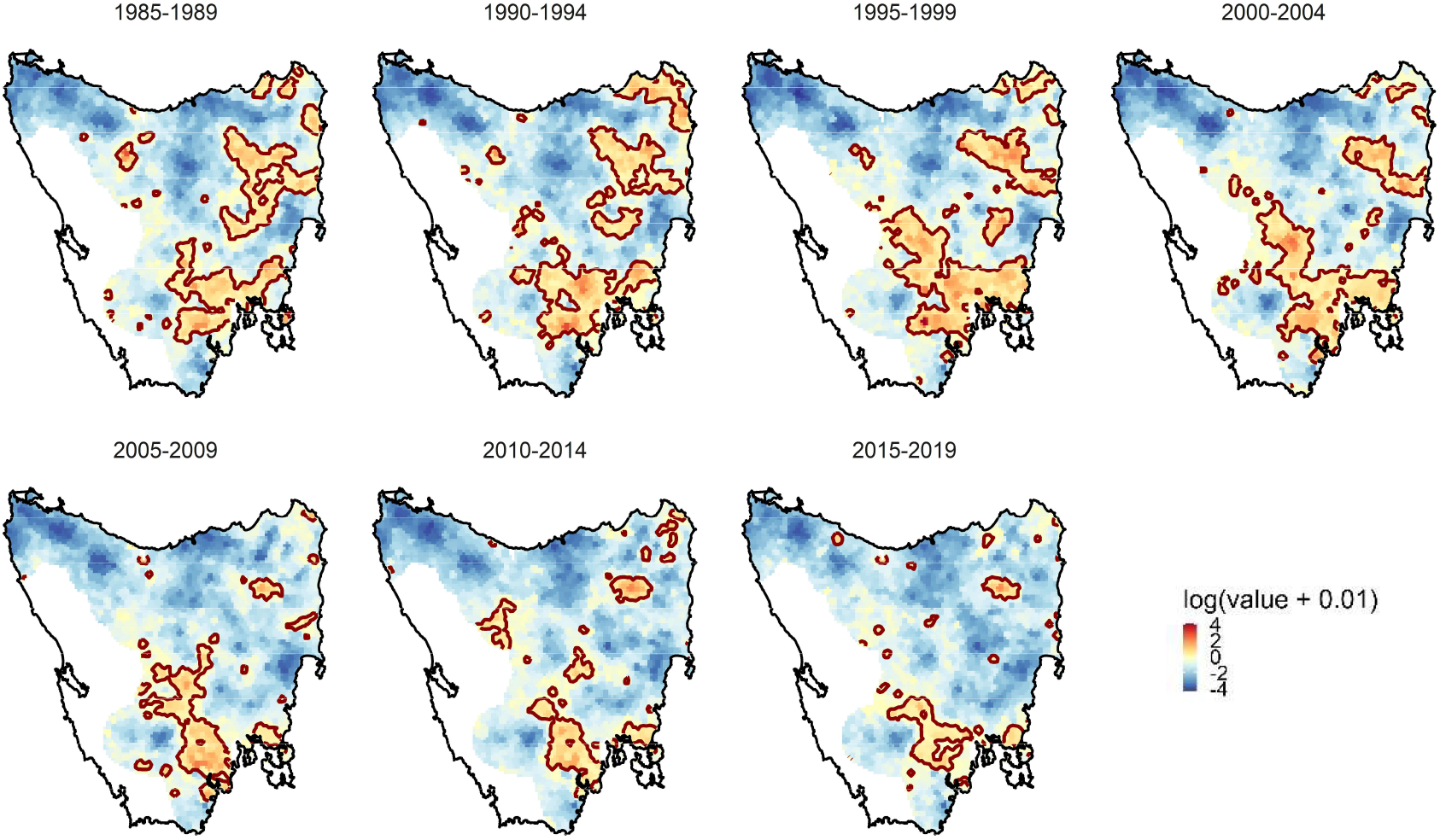
The model-estimated relative abundance (log-transformed for visualisation purposes) of eastern quolls based on a spatiotemporal autoregressive model of spotlight detections. The red contour denotes the model-estimated area of at least one eastern quoll per spotlight transect.

**Figure 3:**
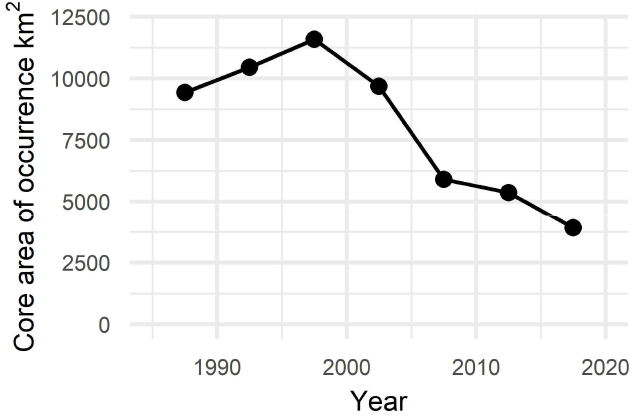
The model-estimated decline in the area occupied by high densities of eastern quolls (defined as a model-estimate of more than one quoll per transect, shown by the red line in Fig 2).

There was strong support for the occurrence of change points that differed among regions (Table 1). The best-performing model for the North, where peak densities were lower than in more southerly populations, indicates a change-point in the early-to-mid 1990s (Fig 2 & 4). In the East and Northeast regions, where initial abundances were among the highest, a distinct step-change occurred in the early 2000’s (Fig 4). In the South region, the population underwent a more recent step-change around 2009. There was no evidence of population change in the Central region.

**Table 1:** Model selection table for the analyses of the regional timing of change points. We selected the best-performing model using the Estimated Log-Predictive Density (ELPD), a leave-one-out cross validation metric, with negative ΔELPD indicating lower predictive accuracy. The standard error refers to the standard error for ΔELPD. ‘Chang-point’ indicates that a model contained a disjoined change-point, ‘variance’ indicates that the segments had difference variance, and ‘null’ indicates that a model did not contain a change-point.

| Model structure | ΔELPD | Std. Error |
| --- | --- | --- |
| Tasmania |  |  |
| ~ change-point + variance | 0 | 0 |
| ~ change-point | -0.32 | 1.16 |
| ~ null | -9.7 | 3.58 |
| North |  |  |
| ~ change-point + variance | 0 | 0 |
| ~ change-point | -7.76 | 6.99 |
| ~ null | -15.25 | 9.99 |
| Northeast |  |  |
| ~ change-point | 0 | 0 |
| ~ change-point + variance | -2.22 | 1.67 |
| ~ null | -3.07 | 2.98 |
| Central |  |  |
| ~ null | 0 | 0 |
| ~ change-point + variance | -0.13 | 2.84 |
| ~ change-point | -0.78 | 0.32 |
| East |  |  |
| ~ change-point + variance | 0 | 0 |
| ~ change-point | -1.48 | 2.77 |
| ~ null | -10.17 | 4.42 |
| South |  |  |
| ~ change-point + variance | 0 | 0 |
| ~ null | -6.03 | 2.5 |
| ~ change-point | -6.24 | 2.63 |

**Figure 4:**
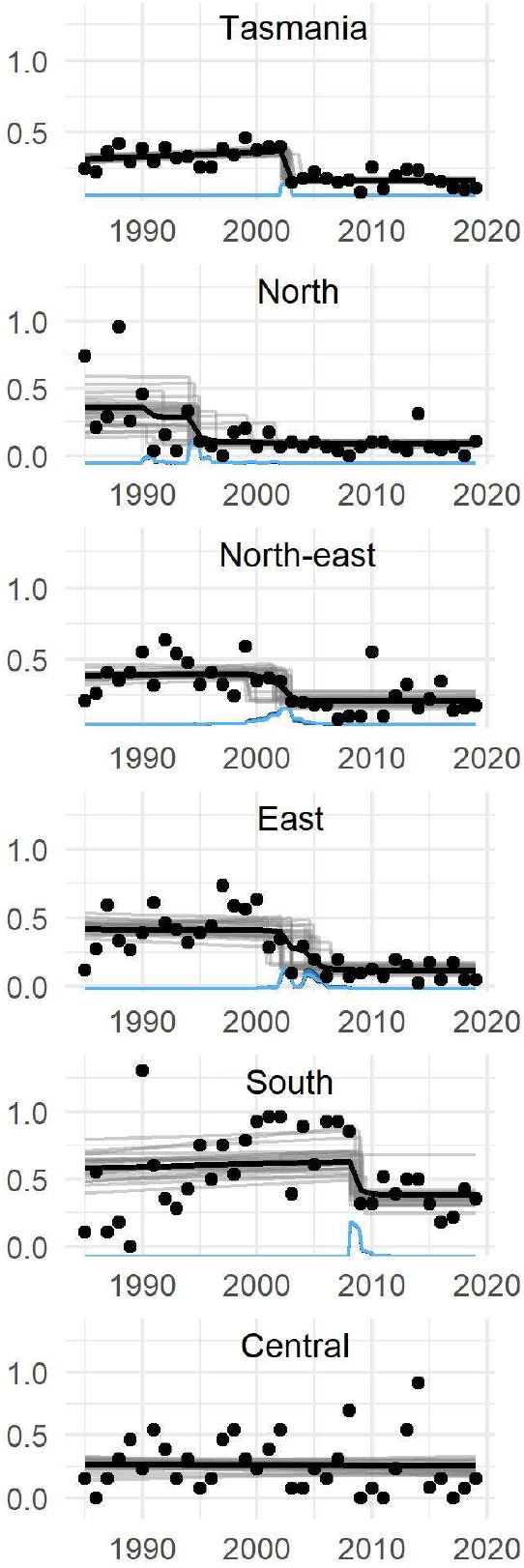
Tasmania-wide and regional change-point models of eastern quoll spotlight detections. Points show the mean number of quolls detected per transect in a year. The black line shows the mean estimate for the best-performing model, with grey lines showing a random subset of model runs. The blue density distribution shows the model-estimated change point. Region names correspond with Fig 1, and are ordered beginning with Tasmania as a whole, and then in ascending order of the estimated change points. The regional models indicate that populations in the North began declining in the mid-1990s, populations in the Northeast and East regions declined in the early 2000s (consistent with the unfavourable weather hypothesis), while populations in the South experienced a more recent decline around 2009.

## Discussion

This study adds to understanding of recent changes in the distribution and abundance of eastern quolls in Tasmania. Using an updated analysis of long-term monitoring data, we confirm earlier conclusions that the Tasmanian population of eastern quolls declined severely during the early 2000s (Fancourt *et al*. 2013; Fancourt *et al*. 2015a). We further show that this range-wide decline was composed of regional declines that occurred at different times, occurring first in the North region and most recently in the South. The general decline of the Tasmania-wide population has continued into recent years with little or no change in trend. The cumulative effect of ongoing declines has been very significant: the total abundance and distribution of eastern quolls in Tasmania has evidently been reduced by more than 60% over the last two decades. The species is extinct on mainland Australia, but until recently it was widespread and abundant in Tasmania and was therefore considered to be safe from complete extinction. We can no longer assume that the existence of eastern quolls in Tasmania ensures its long-term survival.

As well as confirming the general decline of eastern quolls in Tasmania, our results provide new understanding of the pattern of decline. Perhaps most concerning is new evidence of a more recent step-change around 2009 in the South region, where the species was thought to be most secure. We also present new evidence that populations in the North region began declining up to ten years before the onset of decline in the South and East. Pre-decline abundances were lower in the North than in the South and East regions, and the northerly parts of Tasmania have generally lower climatic suitability for the species (Fancourt *et al*. 2015a). Therefore, this geographic pattern of decline could suggest that low-density populations at the periphery of the species’ environmental niche are more susceptible to decline than populations in the core of the niche (Lomolino & Channell 1995). Nonetheless, the ultimate magnitude of reduction in distribution and abundance has been similar in the northern and southern regions. In the Central region, which lies on the cool western margin of the species’ distribution in Tasmania, abundance has been consistently low and there were no indications of a recent change in abundance. A caveat for the central region is that there are fewer spotlight transects, so our power to detect change is lower. Eastern quolls are present at high density on Bruny Island in the far southeast. Bruny Island is not included in the state-wide monitoring program that produced the data analysed here, but anecdotal evidence and unpublished field studies (Cyril Scomparin *pers. comm*.) suggest the species remains abundant there.

It is not clear what has caused the reduction of abundance and distribution of eastern quolls in Tasmania. Fancourt *et al*.’s (2015a) study, which revealed a fall in abundance in the early 2000s, attributed this change to a period of unsuitable weather. The climatic niche of the eastern quoll is characterised by cool dry winters. Between 2000 and 2003, a series of unusually warm and wet winters caused a large temporary contraction in the total area of Tasmania providing that climatic niche. The sharp decline in abundance detected by Fancourt et al (2015a), and confirmed by our analysis, coincided with this event. Why these changed conditions reduced the abundance of eastern quolls is unknown, but Fancourt *et al*. (2018) suggested that effects on the phenology of key invertebrate or vertebrate prey might have been involved. Favourable weather conditions had returned to most of the species’ range by 2005, but abundance did not recover, presumably because some other factor prevented this.

Our results suggest that the causes of decline may be more complex than proposed by Fancourt *et al*. (2015), for two reasons. First, we show that in the North of Tasmania, eastern quoll abundance began falling in the early 1990s, when Fancourt *et al*.’s (2015) analysis suggests weather conditions were generally favourable for the species throughout its range. It remains plausible that changed weather conditions triggered later declines in the East and Northeast, but it is difficult to make that case for the earlier declines in the north. Possibly, some localised weather event triggered those early declines, during a period when eastern quoll populations in the South and East were increasing (as also shown by our analysis), but at present it is not clear what that event might have been. Second, while the weather anomaly of the early 2000s may be able to account for the onset of decline in the core of the species’ range in the Northeast and East, it cannot explain why declines have continued throughout Tasmania since that time. Other causes must be responsible for this, but at present it is not clear what those factors might be, or which of several plausible factors is most important.

A leading candidate is predation by the feral cat (*Felis catus)*. The eastern quoll is among the Australian native mammal species with the highest susceptibility to invasive predators, including the feral cat (Radford *et al*. 2018). Feral cats are widespread in Tasmania, and their densities are highest in the agricultural regions that form a substantial part of the former distribution of eastern quolls (Hollings *et al*. 2014; Hamer *et al*. 2021). Predation by cats, particularly of juvenile quolls, may hold low-density quoll populations in a “predator pit”, and explain why populations that declined during temporary periods of unfavourable weather did not subsequently recover (Fancourt *et al*. 2015b). Elsewhere in Australia, the northern quoll has suffered substantial range contractions to topographically complex environments, presumably where predation from introduced predators poses a lower threat (Moore *et al*. 2019).

Densities of feral cats have increased across much of eastern Tasmania following declines of the Tasmanian devil due to disease (Hollings *et al*. 2014; Hollings *et al*. 2016; Cunningham *et al*. 2020). This disease emerged in northeast Tasmania in the mid-1980s (Patton *et al*. 2020) with substantial population declines commencing in the mid-1990s (Hawkins *et al*. 2006; Cunningham *et al*. 2021). The increasing abundance of feral cats could help to explain the continuing declines of eastern quolls across most of their distributional range. Furthermore, the effects of cats may interact with changes to the structure of low, dense vegetation that provides refuge and escape possibilities (as shown for small mammals in northern Australia; McGregor *et al*. 2014; McGregor *et al*. 2015; Leahy *et al*. 2016). The role of feral cats and vegetation structure in suppressing eastern quolls could be experimentally tested at large scale using translocations of quolls into areas where quolls persist at low densities or have recently disappeared, combined with supplementation of vegetative cover, artificial refuges and short to medium-term control of feral cats. Any investigation of the causes of continuing decline of eastern quolls should also consider possible effects of food availability due to changes in land use and climate.

When the eastern quoll was listed as Endangered under the Commonwealth Environment Protection and Biodiversity Conservation Act 1999, the Threatened Species Scientific Committee (2015) prepared conservation advice and recommended recovery actions. Since publication of this advice, most effort has been directed towards ex-situ management, including the establishment of a captive breeding program under the Tasmanian Quoll Conservation Program, establishment of insurance populations in fenced sanctuaries on the mainland (e.g. Wilson *et al*. 2020), and reintroduction to the wild on the Australian mainland (Robinson *et al*. 2020). Some consideration has also been given to establishing the species on offshore islands (Barlow *et al*. 2021). However, there has been limited work towards halting and reversing declines within their extant Tasmanian distribution, except for plans to manage feral cats on Bruny Island.

In light of the continuing decline of wild populations of eastern quolls, we recommend additional focus on safeguarding populations in Tasmania. First, a critical review of the state-wide monitoring of the species is required. The spotlighting transects provide an invaluable long-term dataset but were designed to monitor large game species (Hocking & Driessen 1992). Supplementing or testing this dataset against other methods would give greater confidence in the state of wild populations and the effectiveness of management actions. Second, preliminary results from a recent pilot study in the Tasmanian central highlands support supplementing wild populations with captive-bred animals (Hamer *et al*. 2022). If this produces a sustained increase in the abundance of wild populations, extending these trials to lowland regions where declines have been most severe could be valuable. Although there is urgency to act while quolls still occupy some parts of Tasmania in moderate abundance, recovery actions need to fit within an experimental and adaptive management framework. Such a framework that monitors the fates of released individuals can provide valuable information on the threats and major causes of mortality at release sites (e.g. Robinson *et al*. 2020) and therefore help to test hypotheses about the factors responsible for the original and ongoing population declines.

In conclusion, declines in eastern quoll populations in Tasmania have continued over the last two decades. The extinction of this species in the wild is a possibility requiring urgent investment and attention. Ensuring that wild quolls persist in the wild should be given high priority, and conservation actions should be embedded within an experimental framework to diagnose current drivers of population decline.

## Acknowledgements

We thank Michael Driessen, Greg Hocking and the Department of Natural Resources and Environment Tasmania for providing the spotlight dataset.

## Data availability

The spotlight data used in this paper were used under agreement with the Department of Natural Resources and Environment Tasmania (formerly Department of Primary Industries, Parks, Water and Environment. Please contact them to obtain the data.

## Conflicts of Interest

The authors declare no conflicts of interest.

## Declaration of Funding

This research did not receive any specific funding.

## Notes

### Competing Interest Statement

The authors have declared no competing interest.

