## Supplementary Material for "Continuing decline of the eastern quoll in Tasmania"

**Table S1:** Summary of the number of spotlight transects surveyed in each region of Tasmania from 1985-2019. Regions are as follows: C = central, N = north, NE = north-east; NW = north-west; E = east; S = south. The maximum variation in the number of transects surveyed in each region (i.e. maximum – minimum number of transects) is as follows: Central =1; North = 7;  North-east = 8; North-west = 0; East = 9; and South = 19.

| Year | C | N | NE | NW | E | S | Total |
|---|---|---|---|---|---|---|---|
| 1985 | 13 | 23 | 42 | 12 | 33 | 9 | **132** |
| 1986 | 13 | 24 | 42 | 12 | 33 | 9 | **133** |
| 1987 | 13 | 24 | 44 | 12 | 39 | 9 | **141** |
| 1988 | 13 | 23 | 45 | 12 | 39 | 11 | **143** |
| 1989 | 13 | 23 | 44 | 12 | 34 | 9 | **135** |
| 1990 | 13 | 24 | 47 | 12 | 41 | 13 | **150** |
| 1991 | 13 | 27 | 47 | 12 | 41 | 20 | **160** |
| 1992 | 13 | 26 | 47 | 12 | 41 | 28 | **167** |
| 1993 | 13 | 27 | 48 | 12 | 41 | 28 | **169** |
| 1994 | 13 | 27 | 48 | 12 | 41 | 28 | **169** |
| 1995 | 13 | 27 | 49 | 12 | 41 | 28 | **170** |
| 1996 | 13 | 28 | 49 | 12 | 41 | 28 | **171** |
| 1997 | 13 | 28 | 49 | 12 | 41 | 28 | **171** |
| 1998 | 13 | 29 | 49 | 12 | 41 | 28 | **172** |
| 1999 | 13 | 29 | 49 | 12 | 41 | 28 | **172** |
| 2000 | 13 | 30 | 49 | 12 | 41 | 28 | **173** |
| 2001 | 13 | 29 | 49 | 12 | 42 | 28 | **173** |
| 2002 | 13 | 29 | 49 | 12 | 41 | 28 | **172** |
| 2003 | 13 | 29 | 49 | 12 | 42 | 28 | **173** |
| 2004 | 13 | 29 | 50 | 12 | 41 | 28 | **173** |
| 2005 | 13 | 29 | 49 | 12 | 41 | 28 | **172** |
| 2006 | 13 | 29 | 49 | 12 | 41 | 28 | **172** |
| 2007 | 13 | 29 | 49 | 12 | 41 | 28 | **172** |
| 2008 | 13 | 28 | 48 | 12 | 41 | 28 | **170** |
| 2009 | 13 | 29 | 49 | 12 | 41 | 28 | **172** |
| 2010 | 13 | 29 | 49 | 12 | 40 | 28 | **171** |
| 2011 | 13 | 29 | 49 | 12 | 41 | 27 | **171** |
| 2012 | 13 | 29 | 49 | 12 | 41 | 28 | **172** |
| 2013 | 13 | 29 | 49 | 12 | 40 | 28 | **171** |
| 2014 | 12 | 29 | 49 | 12 | 41 | 28 | **171** |
| 2015 | 12 | 29 | 49 | 12 | 41 | 28 | **171** |
| 2016 | 13 | 25 | 49 | 12 | 41 | 28 | **168** |
| 2017 | 13 | 29 | 49 | 12 | 41 | 28 | **172** |
| 2018 | 13 | 29 | 49 | 12 | 41 | 28 | **172** |
| 2019 | 13 | 28 | 50 | 12 | 41 | 28 | **172** |
| Total | | | | | | | **5,788** |

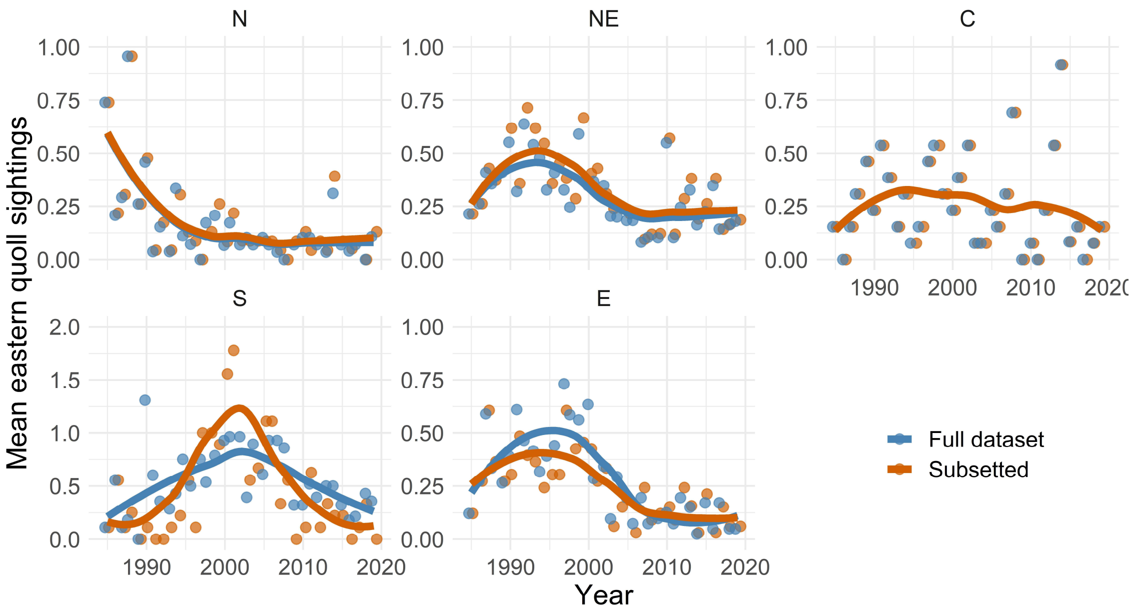


**Figure S1. Visual comparison of the effect of spotlight surveys that were added part-way through the sampling period.** The number of transects surveyed each year increased from approximately 130 in the 1980s to approximately 170 in the 1990s (Table S1). The data points show the mean number of eastern quoll detections in each year, and the line shows a Loess smooth. Blue represents the entire dataset, consisting of 5,788 transect surveys, whereas orange shows only sites that were consistently surveyed since the beginning of data collection in 1985. In general, the effect of adding new transects had a negligible effect on the trends. The most notable effect was in the South region, where the number of transects increased from only nine in the 1980s to 28 in the 1990s (Table S1). Note that we used a different y-axis scale for the South region so that differences in each graph are more easily discernible.
